# Normal aging affects unconstrained three-dimensional reaching against gravity with reduced vertical precision and increased co-contraction

**DOI:** 10.1101/2020.12.03.410001

**Authors:** George F. Wittenberg, Jing Tian, Nick Kortzorg, Lore Wyers, Florian Van Halewyck, Matthieu P. Boisgontier, Oron Levin, Stephan P. Swinnen, Ilse Jonkers

## Abstract

Reaching for an object in space forms the basis for many activities of daily living and is important in rehabilitation after stroke and in other neurological and orthopedic conditions. It has been the object of motor control and neuroscience research for over a century, but studies often constrain movement to eliminate the effect of gravity or reduce the degrees of freedom. In some studies, aging has been shown to reduce target accuracy, with a mechanism suggested to be impaired corrective movements. We sought first to explore the changes in control of shoulder and elbow joint movements that occur with aging during performance of reaching movements to different target heights with the normal effects of gravity, unconstrained hand movement, and stable target locations. Three-dimensional kinematic data and electromyography were collected in 14 young (25±6 years) and 10 older adults (68±3 years) during second-long reaches to three targets aligned vertically in front of the participants. Older adults took longer to initiate a movement than the young adults and were more variable and inaccurate in their initial and final movements. Target height had greater effect on trajectory curvature variability in older than young adults, with angle variability relative to target position being greater in older adults around the time of peak speed. There were significant age-related differences in use of the multiple degrees of freedom of the upper extremity, with less variability in shoulder abduction in the older group. Muscle activation patterns were similar, except for a higher biceps-triceps co-contraction and tonic levels of some proximal muscle activation. The path length of movements was not affected by age. These results show an age-related deficit in the motor planning and online correction of reaching movements against a predictable force (i.e., gravity). These results will facilitate interpretation of our forthcoming study of transcranial magnetic stimulation effects on the same task in these two populations, and is relevant to any study that seeks to measure the effect of pathological processes on upper extremity motor performance in the elderly.

## 1. Introduction

Reaching for an object in space is a movement pattern that forms a basis for many activities of daily living and has long been a subject of human movement science (Elliott et al., 2010). In many studies, reaching is simplified by restricting it to a two-dimensional plane with antigravity support of the arm. This reductionist approach has many benefits, but leaves open the question of whether there is any special consideration to the problem of countering the varying gravitational torques that occur when a multijoint limb is lifted and reaches away from the body. The problem is particularly relevant in neurorehabilitation, where people develop inability to reach against gravity after stroke, and where usual daily activities involve constant compensation for gravitational forces and sometimes harnessing of gravitational forces for intended movements. Before studying reaching against gravity in neurologically impaired patients, normative data from healthy older adults is required. Even in normal aging, there is loss of muscle mass (Moulias et al., 1999;Vandervoort, 2002;Prior et al., 2016) and potentially compensatory increases in brain activity associated with movement (Heuninckx et al., 2005;Heuninckx et al., 2008;Goble et al., 2010). Such data would provide a basis to assess and improve numerous interventions in neurologically impaired patients involving movements with compensation for gravity (Prange et al., 2009;Moubarak et al., 2010;Bastiaens et al., 2011;Grimm et al., 2016).

For the bulk of aging studies, reaching movements have been performed in the horizontal plane, often supported (Przybyla et al., 2011;Coats et al., 2016). In studies in which there was no limb support, reaching movement in the horizontal plane showed higher end-point error and end-point variability in older adults (Poston et al., 2013), and age-related differences in the relative distribution of ballistic and corrective movements (Poston et al., 2009) or ability to learn optimal speed-accuracy tradeoffs (Welsh et al., 2007). Other studies using unsupported reaching in three-dimensional (3-D) space have demonstrated that the end-point spatial variability of corrective movements in response to target displacement during reaching was affected by aging (Kimura et al., 2015). What is not known is the extent to which aging impacts kinematics and muscle activations in a 3-D reaching task against gravity. Such information is necessary to better understand the underlying mechanisms of reaching against gravity in aging and to improve clinical practice. Previously, the laboratory in which this work was performed had shown that reaching against gravity affected the joint coordination strategy when compared to planar movements in young adults (Vandenberghe et al., 2010). Shoulder activation led elbow activation in time, but an elbow control strategy was used to adjust to target height. That study involved restriction of wrist motion, a common strategy to reduce the degrees of freedom, but one that introduces an unrealistic element.

Here, we sought to explore the changes in control that occur with aging in the most naturalistic model we could design and still coordinate with kinematic and electromyographic measures, with the possibility of regional brain stimulation (with data to be presented in the future). In older adults, the ability to reach against gravity may be affected by loss of general muscle bulk (Prior et al., 2016). To the best of our knowledge, this issue has not been studied yet. We investigated the effects of healthy aging and target location on kinematics and muscle activity in a 3-D reaching task against gravity. In terms of kinematics, we hypothesized that 1) older adults would be slower and more variable than young adults in the vertical plane due to poorer integration of predicted effect of gravity on limb movements, but 2) with similar accuracy due to the absence of time constraints on movements (Fitts law) (Boisgontier and Nougier, 2013). In terms of muscle activity, we hypothesized that older adults would show higher levels of cocontraction to improve accuracy (Gribble et al., 2003) and counteract the increased endpoint instability previously described. Multiple vertical targets were used to provide a challenge to motor planning that would not be present with a single target. Finally, the main motivation for collecting and analyzing these motor performance data was to provide a basis for the effects of non-invasive stimulation of cortical areas on performance.

## 2. Material and Methods

### 2.1. Participants

Fourteen young (5 men and 9 women, mean age 25.4 ± 5.9 SD) and 10 older adults (7 men and 3 women, mean age 67.6 ± 3.2) participated in the study. All participants reported good health, with no history of neurological diseases, and normal or corrected-to-normal vision. An Edinburgh inventory was used to assess the handedness (Oldfield, 1971). The study included only right-handed participants. Before any data collection procedures, a written informed consent was signed in agreement with the local ethical committee (1964).

### 2.2. Experiment setup and procedures

The subject was positioned on an adjustable chair with a backrest. An auto-racing restraint system was used to restrict trunk movement. An adjustable table with a visual stimulus presentation system was placed in front of the subject (Figure 1A.) Three pairs of light emitting diodes (LEDs) represented an upper target (UT), middle target (MT) and lower target (LT). Both the LT and the UT were vertically separated 15 cm from the MT. The height and horizontal position of the MT were aligned to the right shoulder. The distance to the MT was set to 5 cm less than a fully extended reach to the MT. The starting position was marked on the table and in line with the targets, 3 cm lower than the LT, 15 cm from the target board. Subjects were asked to find a comfortable sitting position with right upper arm in vertical and adducted position and elbow flexed approximately 90 degrees. The forearm was prone with the hand resting on the table and the tip of index finger on the starting position.

**Figure 1.**
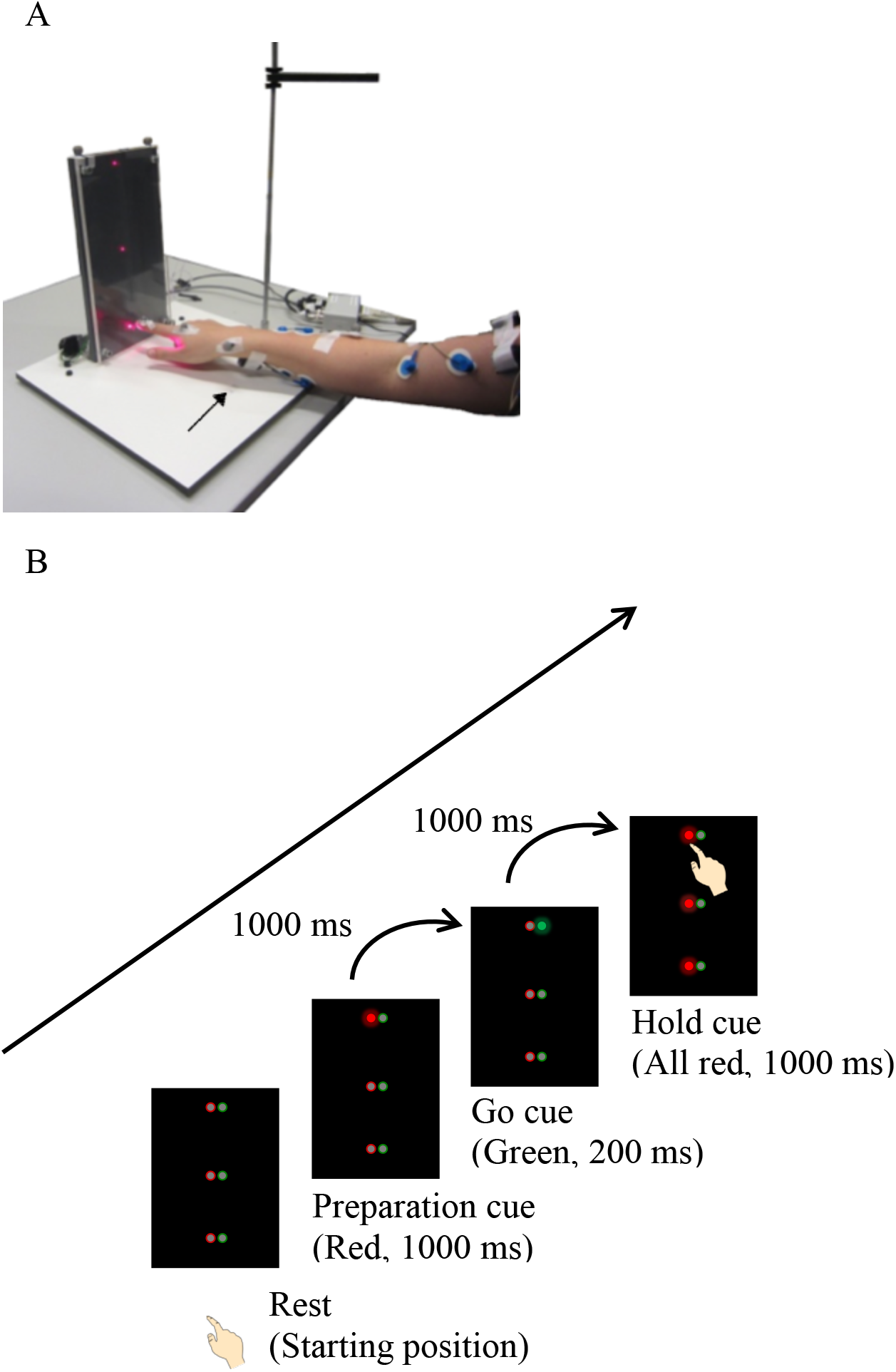
(A) Physical setup with photo of subject reaching to lower target with EMG electrodes and reflective markers attached (see Methods for details). The arrow indicated the starting position for the fingertip. (B) Sequence of events for a single reach.

Each target of the visual stimulus presentation system contained one red (left) and one green (right) LED, separated 1 cm from each other (Figure 1B). Participants were instructed to start at rest with the tip of their index finger on the starting position. Relaxation prior to movement initiation was stressed explicitly. First, one red light illuminated for 1 s as a preparation cue, indicating which target would be the goal of the reaching movement. After an additional 1 s delay, a green light flashed for 200 ms as a go cue. The subject performed a smooth reaching movement to the remembered red target. One second after the go cue, all of the red lights illuminated. Participants were instructed to get to the target approximately 1 s after the go cue by attempting to match target contact with this last signal and keep their finger on the target until all the red lights extinguished after an additional second. The hand returned to the starting position to end the reaching cycle. The room light was dimmed, but visual feedback of LED target and the hand position was still available.

A fixed pseudorandom sequence of 24 reaching movements ran automatically with 18 trials related to the TMS protocol and 6 trials without stimulation. In those latter trials, three targets height conditions were tested with each condition repeated twice. The participants practiced a sequence of 24 reaches prior to the actual measurements and then completed 2 to 3 blocks (depending on overall experimental time available) of 24 reaches for each of four different stimulation locations (but with 6 reaches in which no stimulation was given). Here, we only analyzed trials that were performed in the no-stimulation condition, with a paper on the effects of stimulation to follow.

### 2.3. Kinematic recordings

Kinematic data from the trunk and right upper limb were collected at 100 Hz using a 3D motion analysis system (Vicon, Oxford Metrics, UK). Reflective markers were attached at cervical vertebra 7, thoracic vertebra 10, sternoclavicular joint, xiphoid process, acromioclavicular joint, medial and lateral epicondyle of humerus, radial and ulnar styloid process, metacarpophalangeal joint and distal phalanx of the index finger. Additionally, two clusters of markers were used: one on the humerus and one on the forearm. The position of the target board was also recorded using the reflective markers. 3D marker location and EMG signals from the 9 muscles of the right upper limb (see description below) were collected synchronously. Static calibration trials were collected prior to the dynamic trials.

### 2.4. EMG recordings

Electromyography (EMG) surface electrodes (Red Dot, 3M, Diegem, Belgium) were attached to the skin overlying the following muscles: pectoralis superior (PEC), trapezius pars descendens (TRP), anterior deltoid (DLA), medial deltoid (DLM), posterior deltoid (DLP), biceps brachii (BIC), triceps long head (TRI), flexor carpi ulnaris (FCU), and extensor digitorum communis (EDC). Signals were amplified and collected at 1000 Hz using the ZeroWire wireless EMG system (Aurion, Milan, Italy).

### 2.5. Data analysis

Final data analysis was performed for 10 young and 8 older participants with high-quality kinematic and EMG recordings. (First 4 young participants were excluded because they were tested to optimize the experimental design and one older participant was excluded due to missing shoulder marker and the other due to failure to follow the instruction). For endpoint kinematic and EMG analysis, each young subject contributed between 11 and 23 trials and older participants between 13 and 23 trials for each target location. For joint kinematic analysis, data from one young participant were further excluded due to missing markers on the trunk in some trials.

#### 2.5.1. Kinematic data analysis

##### 2.5.1.1. Pre-processing

The recorded 3D positions of the reflective markers were reconstructed and labeled in Nexus (Vicon). The reflective marker positioned on distal phalanx of index finger was used for the calculation of endpoint kinematic variables. The index finger trajectory was transformed into the reference frame of the target board, with the origin at the upper left corner of the board, the x-axis perpendicular to the board, the y-axis horizontal (parallel to the upper edge of the board) and the z-axis vertical. Upper body joint angles were calculated as the following: anteflexion (ShFlx), abduction (ShAbd) and internal rotation (ShRot) of the humerus with respect to the trunk, elbow flexion (ElFlx), pronation (WrPrn) of the forearm with respect to the humerus, and ulnar deviation (WrDev) and extension of the hand (WrExt) with respect to the forearm.

Kinematic data were further processed using custom software developed in MATLAB™ (The Mathworks, Natick, MA). A low-pass fourth-order, zero lag Butterworth filter was applied to kinematic data with a cutoff frequency of 10 Hz. Marker velocities along each of the above axes were calculated by determining the derivative of the position signal. 3D speed of the index finger was calculated as the magnitude of the velocity vector. The initiation and the end of the endpoint movement were determined using a threshold of 5 % of the 3D peak speed of the metacarpal marker. All trials were visually inspected to ensure the accuracy of the automatic procedure. Trials were rejected if the movement started before the go cue or the reaction time was less than 50 ms. Abnormal trials were further excluded using criteria as follows: reaction time differed > 2 SDs from the average, and movement duration > 2 SDs longer than the average. As a result, 13% of the young and 17% of the older participants’ trials were excluded. Shoulder and elbow movement onsets and offsets were defined as described above for the endpoint.

##### 2.5.1.2. Endpoint kinematics

###### Time-related variables

*Reaction time* was calculated as the time interval between the go cue and the initiation of the movement, and *movement time* as the interval between the initiation and the end of movement. *Response time* was defined as the sum of reaction time and movement time. *Peak speed* was calculated for each movement. *Speed skewness,* or asymmetry, was defined as the ratio of the duration of the acceleration phase to the total movement duration. Skewness of 0.5 denotes a symmetric speed profile.

###### Space-related variables

*Path length* was calculated as the sum of subsequent distances between adjacent data points along the movement path. The *index of curvature* was defined as the ratio between the path length and the distance between the position of the finger at the onset and at the end of each movement. *Endpoint precision* was evaluated as the variability (within subject SD) at the end of the movement for each of the three axes separately.

*Spread of paths* was further assessed by calculating the magnitude and angle of the 3D reach vector at different stages of movement. Specifically, early motor planning was assessed by calculating the magnitude and angle of the 3D reach vector at 100 ms after movement onset and peak speed. Online feedback control was assessed by analyzing the reach vectors at 50 ms after peak speed and movement end. The 3D reach vector was defined based on the position of the finger at the onset of movement and at each instant. The magnitude was calculated as the square root of the sum of squares of the reach vector component along the three motion axes. The angle of the reach vector was calculated between the actual 3D movement vector and the straight-line path to target. The variability (within subject SD) of the magnitude and angle was also calculated for the different time points to characterize the control of movement.

##### 2.5.1.3. Joint kinematics

The *joint excursion range* was calculated for three DOFs of the shoulder, one DOF of the elbow and three DOFs of the wrist. The *variability of joint angles* was evaluated as the within subject SD of seven joint angles at the same four instances as for the analysis of the endpoint paths. To compensate for small inter-trial variations of the actual starting position of the arm and of movement duration, we corrected the raw joint angles using a linear regression model with the predictor of seven DOFs of initial arm position and movement duration (Krüger et al., 2011). This correction was performed separately for each subject, each target position and for each of the four instances.

#### 2.5.2. EMG data analysis

Raw EMG signals were band-pass filtered (10–400 Hz) using a fourth-order, zero-lag Butterworth filter and rectified (Bosch et al., 2009). The envelope of the rectified signal was calculated using a moving window with bin size of 3 ms. Maximum voluntary contraction was subjectively more difficult to obtain in older adults. Consequently, to enable comparisons across participants, EMG data for each muscle were first normalized for each subject by dividing by the maximum observed EMG activity for that muscle during the experimental session. EMGs were then time-aligned to movement onset for each trial and averaged. The subsequent analyses were based on averaged EMG data between 100 ms before movement onset and 100 ms after movement end. Results from the DLP and FCU muscles were not reported here, since their signals were close to noise level in some participants.

Onset of muscle activity was determined from averaged data, i.e., the time when the EMG first exceeded the resting baseline (mean of the first 100 ms after the go cue) by at least 3 SDs for a minimum of 10 ms. All EMG onset times were normalized to movement onset. Peak amplitude of normalized EMG was determined from averaged data for each muscle. Tonic EMG levels following movement were determined for each muscle by computing the mean level of EMG activity during a 100-ms period after the movement ended. Note that the measurement of tonic EMG was conducted on the basis of individual trials. To assess the co-contraction of shoulder-elbow muscles, tonic EMG activity of BIC and TRI were averaged.

### 2.6. Statistical analysis

A linear mixed model was performed on each of the dependent variables with age-group and target location as the fixed effect and subject as the random effect (Boisgontier and Cheval, 2016). Significant effects were determined using a likelihood ratio test to compare pairs of models (with and without the particular factor of interest; *p* values are reported along with the corresponding χ^2^ value). If the location effect was significant, *F*-tests on the fixed effects coefficients were applied to examine pairwise differences; reported *p* values were not further corrected. Pearson’s linear correlations were calculated for the co-contraction of BIC and TRI and the endpoint variability. These statistical analyses were conducted using MATLAB™.

## 3. Results

Our reaching tasks required the two movement elements of vertical lift and forward reach. Participants were free to choose any path to reach the target but trunk movement was restricted. During reaching, only minimal movement of the trunk was observed (backward tilt: 1.2 ± 0.6° SD, left tilt: 0.6 ± 1.0° and left rotation: 2.7 ± 1.5°). Figure 2 shows the typical endpoint trajectories and seven joint angles of reach movements to the three different target locations in an example young (A, C) and older adult (B, D). Visual inspection of the reach trajectories in both young and older participants showed that they often curved and changed direction in idiosyncratic ways. Moreover, the finger took a somewhat different path each time to reach the same target. The variation in joint angles of the arm over repetition of movements was rather small. However, the pattern of joint motion during reaching to three different heights varied between participants. Some participants (Fig. 2C) used more abduction and external rotation in the shoulder but less supination and very little flexion in the wrist. However, some participants ((Fig. 2D) used less abduction and external rotation in the shoulder together with relatively greater extent and longer duration of wrist motion. In the following section, we first present the analyses on the kinematic characteristics of endpoint and joint motion.

**Figure 2.**
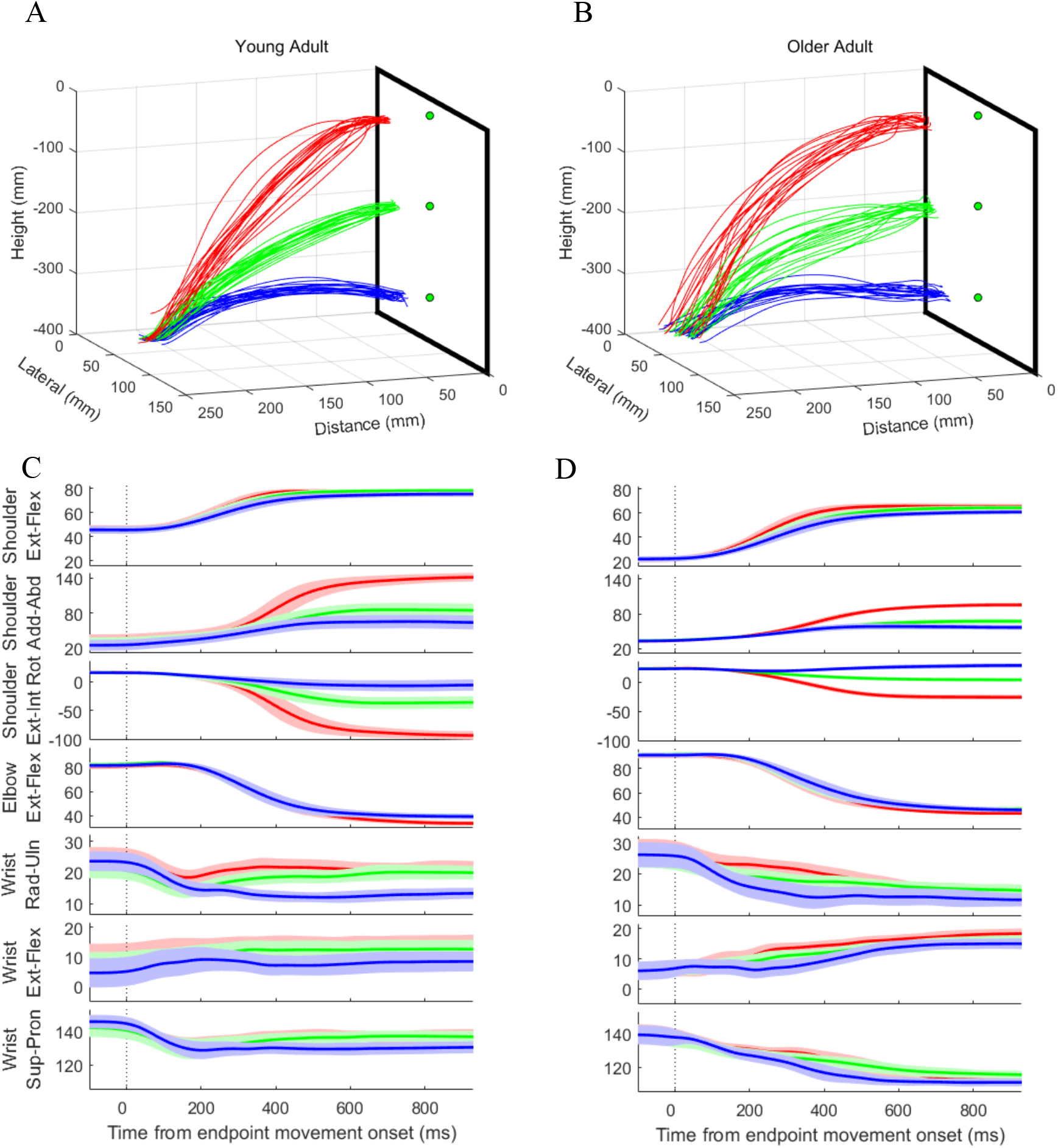
3D view of the endpoint trajectories (A, B) and mean time course of joint angles (C, D) for reaching movements to the three targets heights by a young (A, C) and an older (B, D) adult subject. Each line (A, B) represents a separate trial obtained from a marker positioned on the distal phalanx (endpoint) of the right index finger. The green circles indicate the positions of the targets. Shaded areas (C, D) indicate standard deviation across individual trials. Joint angles are aligned to endpoint movement onset (dotted vertical line, Time = 0 ms).

### 3.1. Endpoint kinematics

#### 3.1.1. Time-related variables

The overall characteristics of time-related kinematics in both groups are shown in Table 1.

**Table 1.** Time-related kinematic variables of endpoint movements in young and older adults.

|  | Young group |  |  | Older group |  |  | Statistics |
| --- | --- | --- | --- | --- | --- | --- | --- |
|  | UT | MT | LT | UT | MT | LT |  |
| Reaction time (ms) | 299<br>(13) | 291<br>(22) | 271<br>(9) | 347<br>(35) | 345<br>(33) | 329<br>(31) | * †† |
| Movement time (ms) | 848<br>(25) | 808<br>(29) | 858<br>(31) | 894<br>(65) | 891<br>(78) | 925<br>(77) | †† |
| Response time (ms) | 1147<br>(31) | 1100<br>(28) | 1130<br>(29) | 1242<br>(77) | 1236<br>(92) | 1254<br>(93) | ns |
| Peak speed (mm/s) | 1060<br>(50) | 771<br>(38) | 575<br>(34) | 1073<br>(71) | 775<br>(48) | 606<br>(40) | ††† |
| Speed skewness | 0.32<br>(0.01) | 0.35<br>(0.01) | 0.35<br>(0.02) | 0.33<br>(0.01) | 0.34<br>(0.01) | 0.37<br>(0.02) | ††† |

Results are presented as mean (SE). Significant age group differences are indicated by \* (if $p < 0.05$ ); target position differences by † (if $p < 0.05$ ), †† (if $p < 0.01$ ), ††† (if $p < 0.001$ ); nonsignificant differences by ns.

##### Reaction Time

Reaction time showed a significant effect of age (χ^2^ (1) = 4.41, *p* = 0.036), with older adults exhibiting longer reaction time than young adults. There was also a significant effect of target location (χ^2^ (2) = 11.7, *p* = 0.003). Pairwise comparisons revealed that in both age groups the reaction time was greater for UT compared with LT (*F*(1,18) = 16.37, *p* < 0.001). No significant interaction between age and target location was found.

##### Movement Time

Movement time showed no effect of age (χ^2^ (1) = 0.23, *p* = 0.635), but a significant effect of target location (χ^2^ (2) = 13.29, *p* = 0.001). Pairwise comparisons revealed that movement times were longer in the LT than MT condition (*F*(1,18) =19.45, *p* < 0.001). No significant interaction between age and target location was found.

##### Response time

Response time was calculated as the time elapsed between the go cue and the end of movement. Response time showed no effect of age (χ^2^ (1) = 0.90, *p* = 0.342) and target location (χ^2^ (2) = 4.75, *p* = 0.093) and no interaction between age and target location. These results are consistent with the enforcement of a response time of ≈ 1s regardless of group and target locations.

##### Peak speed

Peak speed was similar for the young compared with the older adults (χ^2^ (1) = 0.18, *p* = 0.668), but increased with target height (χ^2^ (2) = 52.78, *p* < 0.001). Pairwise comparisons revealed that the peak speed was larger in the UT than LT condition (*F*(1,18) = 317.61, *p* < 0.001), but the difference between any two neighboring targets only showed a trend (MT *vs* LT: *F*(1,18) = 4.36, *p* = 0.051; UT *vs* MT: *F*(1,18) = 3.78, *p* = 0.068).

##### Speed skewness

The skewness of reaching movements in both age groups ranged from 0.32 to 0.37, revealing that participants typically spent proportionally more time after reaching peak speed than before. There was no effect of age (χ^2^ (1) = 0.23, p = 0.634), but a significant effect of target location (χ^2^ (2) = 15.42, *p* < 0.001) for skewness. Pairwise comparison indicated that skewness of reaching movement to UT was lower than to MT and LT (UT *vs* MT: *F*(1,18) = 5.41, *p* = 0.032; UT *vs* LT: *F*(1,18) = 18.01, *p* < 0.001), but there was no significant difference between MT and LT (*F*(1,18) = 2.17, *p* = 0.158).

#### 3.1.2. Space-related variables

##### Path length and variability

The older subjects’ path lengths were similar to those of young subjects (Table 2; χ^2^ (1) = 0.66, *p* = 0.416), but increased with target height (χ^2^ (2) = 95.29, *p* < 0.001). The variability of the path length was also similar between groups (χ^2^ (1) = 0.61, *p* = 0.433) but was not affected by the target location (χ^2^ (2) = 5.02, *p* = 0.081). No significant interaction between age and target location was found for path length and its variability.

**Table 2.** Characteristics of the endpoint trajectories in young and older adults.

|  | Young group |  |  | Older group |  |  | Statistics |
| --- | --- | --- | --- | --- | --- | --- | --- |
|  | UT | MT | LT | UT | MT | LT |  |
| Path length (mm) | 412<br>(7) | 297<br>(9) | 239<br>(12) | 411<br>(5) | 300<br>(6) | 243<br>(7) | ††† |
| Path length variability (mm) | 10<br>(1) | 8<br>(1) | 11<br>(1) | 9<br>(1) | 11<br>(1) | 13<br>(3) | ns |
| Index of curvature (%) | 107.3<br>(0.9) | 109.9<br>(1.5) | 118.8<br>(2.6) | 106.6<br>(1.1) | 109.8<br>(2.2) | 115.4<br>(3.4) | ††† |
| Curvature variability (%) | 2.5<br>(0.3) | 3.0<br>(0.3) | 5.0<br>(0.5) | 2.2<br>(0.3) | 3.6<br>(0.5) | 5.4<br>(1.2) | ††† ‡ |

Results are presented as mean (SE). Significant age group differences are indicated by \* (if $p < 0.05$ ); target position differences by † (if $p < 0.05$ ), †† (if $p < 0.01$ ), ††† (if $p < 0.001$ ); interaction between age and target position by ‡; nonsignificant differences by ns.

##### Index of curvature and variability

Results of the curvature analysis showed that indexes were always greater than 1, indicating that the paths used were never straight (Table 2). There was no effect of age (χ^2^ (1) = 0.51, *p* = 0.477), but a significant effect of target location (χ^2^ (2) = 22.61, *p* < 0.001). Pairwise comparisons showed that the indexes of curvature for the three target heights were significantly different from each other. These results indicated that the amount of deviation from a straight-line path depended on the height of the target. The least deviation was observed for the UT, with progressively greater deviation for the LT. Similarly, no age effect (χ^2^ (1) = 0.11, *p* = 0.746) but an effect of target height (χ^2^ (2) = 21.52, *p* < 0.001) was observed for the variability of curvature; the higher the target the less variable the path curvature was. However, a significant interaction between age and target location was found for curvature variability (χ^2^ (2) = 6.45, *p* = 0.040), indicating that target height had greater effect on the older adults’ curvature variability.

##### Endpoint precision at the end of movement

Endpoint variability on the x-axis (anteroposterior) was similar for the young and older participants (χ^2^ (1) = 0.03, *p* = 0.869; Figure 3A). However, there was a significant effect of target location (χ^2^ (2) = 7.18, *p* = 0.028). Pairwise analyses revealed that endpoint variability on the x-axis was greater for LT compared with UT and MT (UT *vs.* LT: *F*(1,18) = 5.05, *p* = 0.037; MT *vs.* LT: *F*(1,18) = 7.80, *p* = 0.012). For endpoint variability on the y-axis (lateral), the effects of age (χ^2^ (1) = 3.16, *p* = 0.075) and target location (χ^2^ (2) = 2.82, *p* = 0.245) were not significant (data not shown). In contrast, endpoint variability on the z-axis (vertical) was greater for the older participants compared with the young participants (χ^2^ (1) = 5.02, *p* = 0.025; Figure 3B). There was no effect of target location (χ^2^ (2) = 0.87, *p* = 0.648). These results indicated an age-related decline in endpoint precision along the vertical axis regardless of target height, and reduced endpoint precision to the LT along the anteroposterior axes, irrespective of age group.

**Figure 3.**
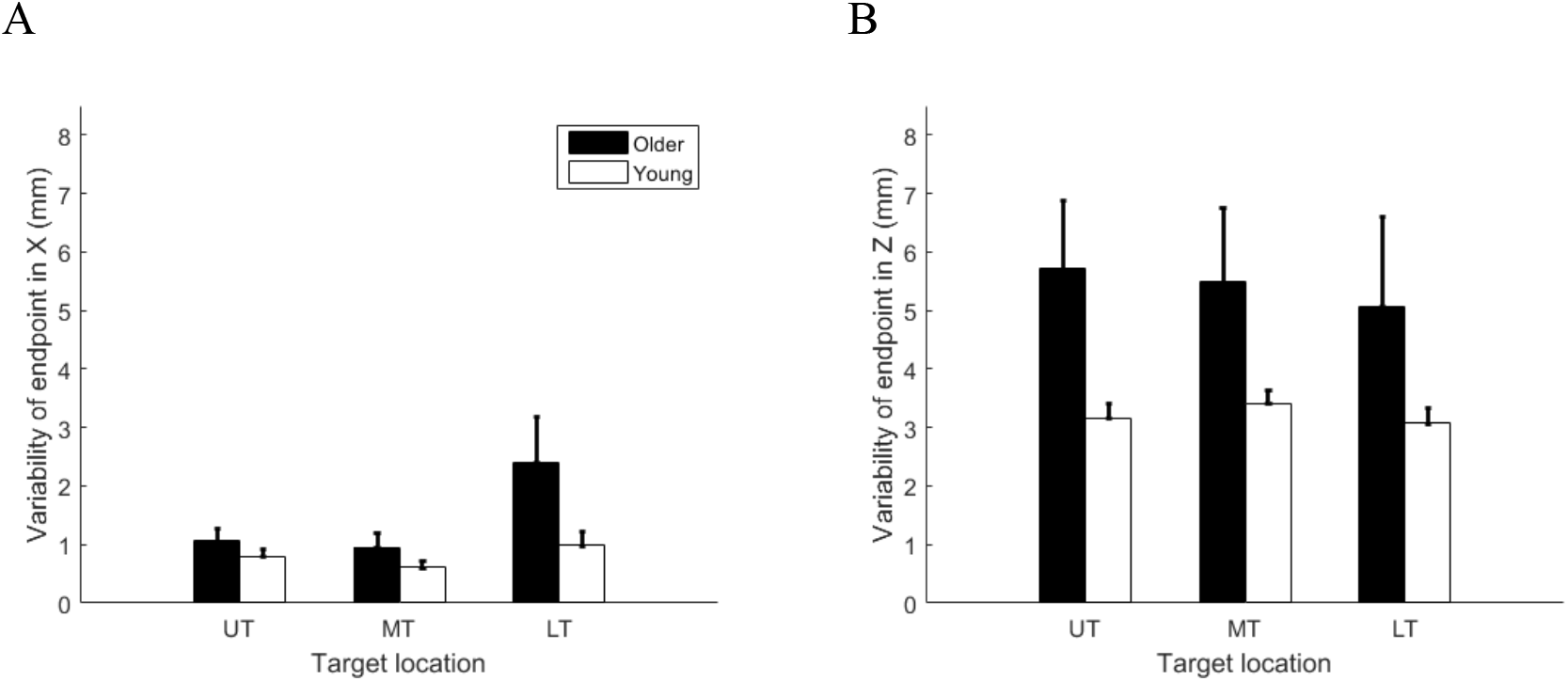
Variability of endpoint on (A) X-axis (anteroposterior) and (B) Z-axes (vertical) for three different target locations in the young and older adults (mean ± SE).

##### Spread of paths at different stages of movement

To examine the effects of age on motor planning and online control during reaching movements, we further assessed magnitude and angle of the reach vector at different stages of movement (Figure 4).

**Figure 4.**
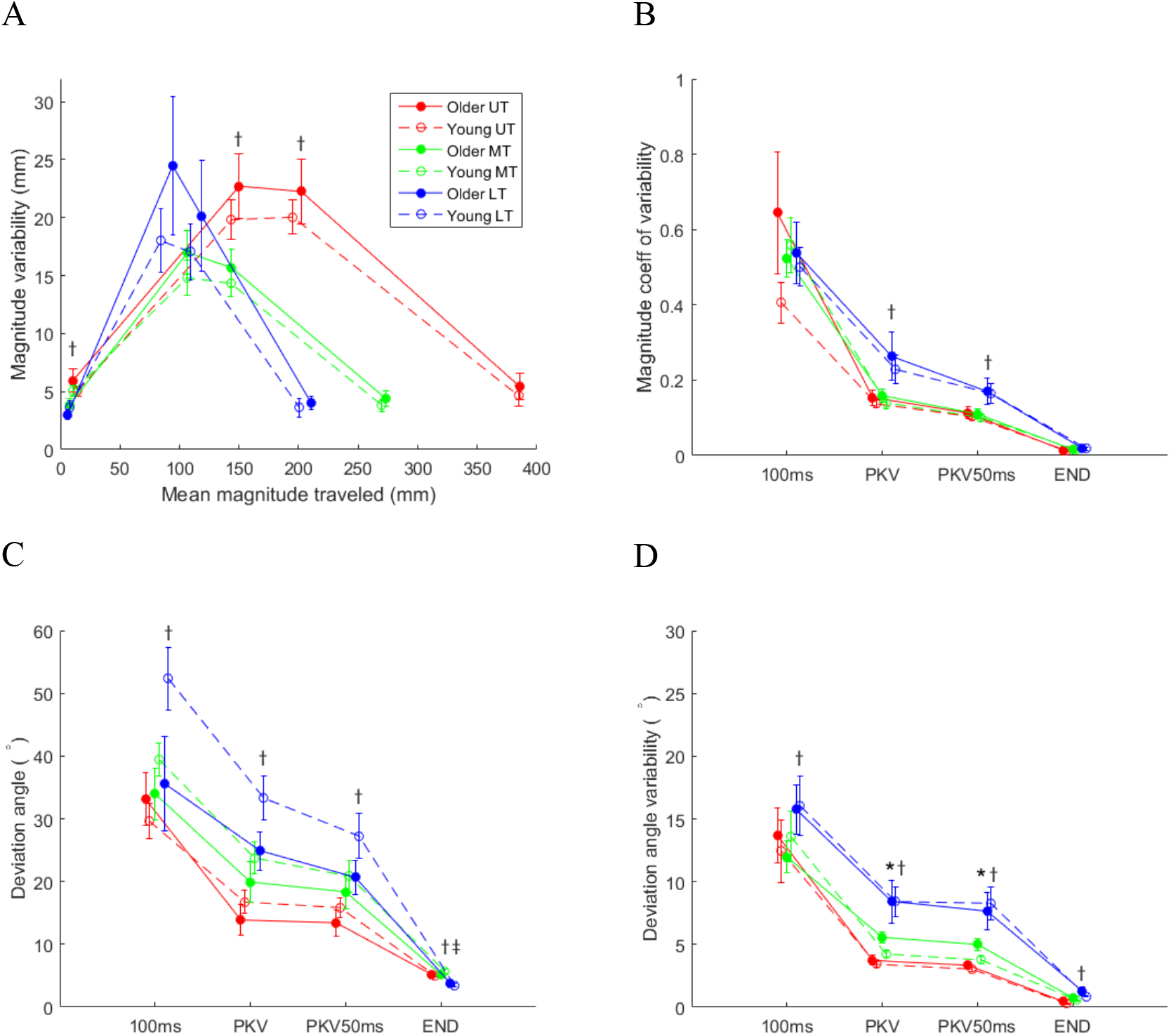
Variability in magnitude of reach vectors vs. mean magnitude of distance traveled (A) and coefficient of variation (B), deviation angle (C) and its variability (D) of reach vectors at 100 ms after movement onset, peak speed (PKV), 50 ms after peak speed (PKV50ms), and movement end (END) for the three different target locations in the young and older subjects (means ± SE). Significant age group differences are indicated by *; target position differences by †; interaction between age and target position by ‡.

In Figure 4A, movement magnitude variability at different time points is plotted as a function of mean magnitude traveled for the three target locations. For all three targets, there were increases in variability up to peak speed but then variability decreased towards the end of the movement. There was no effect of age on magnitude and its variability for all the time points we evaluated (all *p* > 0.390). However, the effects of target locations on magnitude variability were different across the different stages of movement. At 100 ms after movement onset, magnitude variability was lower for LT compared with UT and MT (UT *vs* LT: *F*(1,18) = 14.96, *p* = 0.001; MT *vs* LT: *F*(1,18) = 15.04, *p* = 0.001). At the time of peak speed and 50 ms after peak speed, magnitude variability was lower for the MT than the UT (all *p* < 0.001). There was no significant effect of target locations at the end of movement (χ^2^ (2) = 3.68, *p* = 0.159).

To assess whether the changes in variability were related to the distance traveled, we calculated the coefficient of variation (Figure 4B). If endpoint trajectories were corrected during movement execution, then the coefficient of variation should decrease as the movement unfolds, as we found. But there was no effect of age during any stage of movement (all *p* > 0.382). The effect of target location was found to be significant only at the time of peak speed and 50 ms after peak speed (PKV: χ^2^ (2) = 8.81, *p* = 0.012; PKV50ms: χ^2^ (2) = 10.22, *p* = 0.006). The coefficient of variation was smaller for both UT and MT compared with LT (all *p* < 0.008).

Figure 4C and 4D shows the deviation angles between the reach vector and the target vector and their variability at four time points for three target locations. As the movement unfolded, the deviation angle decreased over time (Figure 4C). In general, reach to LT deviated the most from the intended direction during the early part of the movement. There was no effect of age (all *p* > 0.080), but a significant effect of target location (all *p* < 0.038) on deviation angle for all the time points assessed. The deviation angles for three target locations were significantly different from each other. A significant interaction between age and target location was found only at the end of movement (χ^2^ (2) = 9.64, *p* = 0.008), showing that target height had greater effect on the young participants’ landing direction as compared to the older participants.

The variability of the deviation angle was calculated to further examine the precision of the planning and execution of movement (Figure 4D). There was a significant effect of target location for all four time points (all *p* < 0.016). At 100 ms after movement onset, the angle variability was lower for MT compared with LT (*F*(1,18) = 9.53, *p* = 0.006); at the remaining time points, the variabilities for three target locations were significantly different from each other. The effect of age was found to be significant only at the time of peak speed and 50 ms after peak speed (PKV: χ^2^ (1) = 4.63, *p* = 0.031; PKV50ms: χ^2^ (1) = 3.87, *p* = 0.049), indicating that the angle variability was greater in older participants than young participants around the time of peak speed.

### 3.2. Joint kinematics

#### Joint excursion range

Figure 5 shows the mean joint excursion range as a function of target location for the 7 DOFs in both young and older groups. The shoulder and elbow joints contributed primarily to the reaching movement, whereas the wrist joint typically showed little overall excursion. A significant main effect of age was found in ShFlx (χ2 (1) = 3.88, *p* = 0.049) and ShRot (χ2 (1) = 6.13, *p* = 0.013). The older participants used significantly more shoulder flexion but less external rotation than the young participants. The main effect of target height was found to be significant in all 7 DOFs (ShFlx: χ2 (2) = 29.71, *p* < 0.001; ShAbd: χ2 (2) = 22.45, *p* < 0.001; ShRot: χ2 (2) = 45.04, *p* < 0.001; ElFlx: χ2 (2) = 46.58, *p* < 0.001; WrDev: χ2 (2) = 9.76, *p* = 0.008; WrExt: χ2 (2) = 21.76, *p* < 0.001; WrPrn: χ2 (2) = 7.06, *p* = 0.029). Pairwise comparisons showed that the excursion range in the shoulder’s three DOF for three target heights were significantly different from each other (all *p* < 0.001). The elbow extension was greatest when reaching to UT but smallest to MT (UT *vs* LT: F(1,17) = 37.57, *p* < 0.001; MT *vs* LT: F(1,17) = 13.30, *p* = 0.002; UT *vs* MT: F(1,17) = 196.99, *p* < 0.001). Wrist radial deviation was greater when reaching to LT than UT and MT (UT *vs* LT: F(1,17) = 9.04, *p* = 0.008; MT *vs* LT: F(1,17) = 12.87, *p* = 0.002). Wrist extension increased with target height and a significant interaction between age and target location was found (χ^2^ (2) = 6.84, *p* = 0.033), indicating that target height had greater effect on the older adults’ wrist extension. The forearm became less pronated at the end of the movement and excursion range was only different between UT and MT (UT *vs* MT: F(1,17) = 7.35, *p* = 0.015).

**Figure 5.**
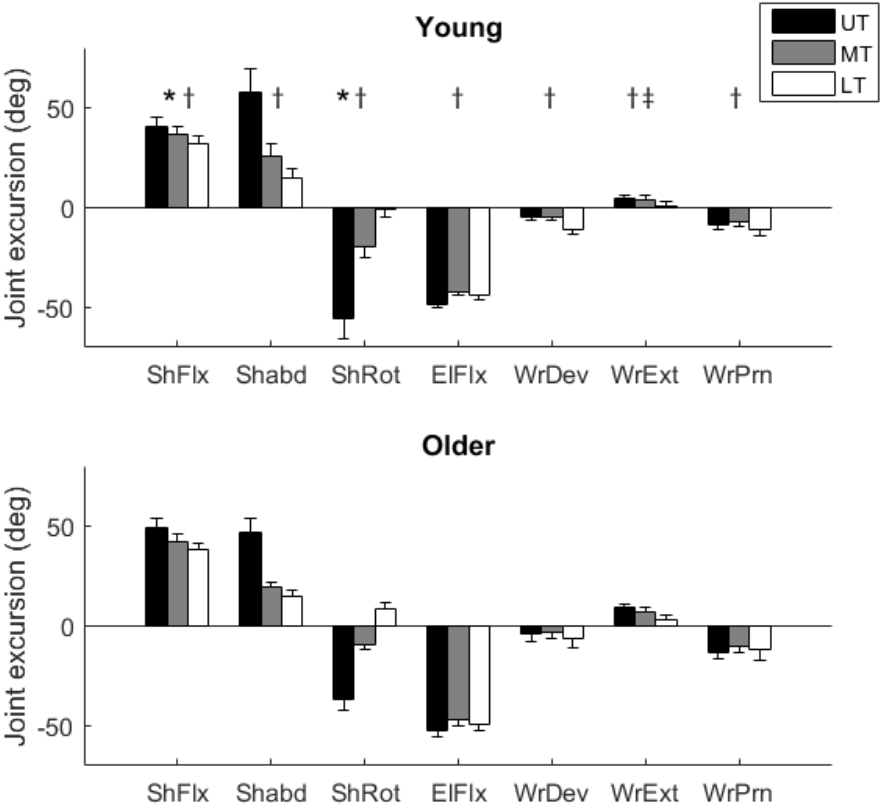
Joint excursion range plotted as a function of target location for 7 DOF in the young and older subjects (means ± SE). Positive angles indicate the following directions: flexion, abduction and internal rotation in shoulder, flexion in elbow, ulnar deviation, extension and internal rotation in wrist. Significant age group differences are indicated by *; target position differences by †; interaction between age and target position by ‡.

#### Joint angle variability at different stages of movement

Figure 6 shows the changes in joint angle variability over time for three target locations. In general, older adults showed a similar temporal evolution of joint angle variability to young adults for each of seven joint angles. For shoulder flexion, elbow extension and wrist extension, the variability showed a similar increase and then decrease pattern with its maximum at around the time of peak speed of the endpoint. For shoulder abduction, shoulder external rotation, wrist radial deviation and wrist internal rotation, the variability increased and then either kept increasing or stabilized at the end of movement depending on the target heights.

**Figure 6.**
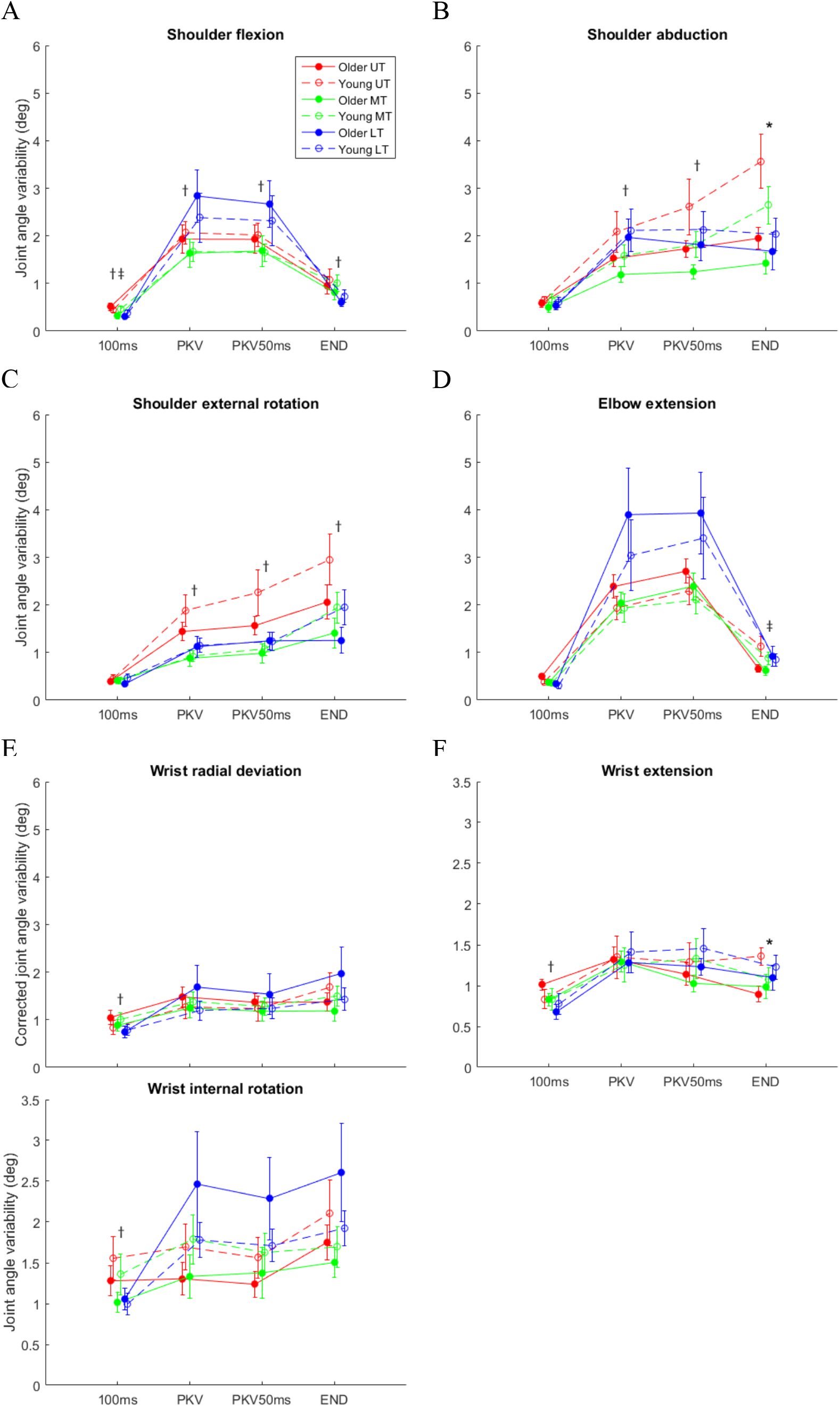
Variability (within-subjects standard deviation) of joint angles for the three different target locations plotted at 100 ms after movement onset, peak speed (PKV), 50 ms after peak speed (PKV50ms), and movement end (END) in the young and older subjects (means ± SE). (A) – (G) present for each single joint angle. Significant age group differences are indicated by *; target position differences by †; interaction between age and target position by ‡.

A significant main effect of age was only found in the variability of shoulder abduction and wrist extension at the end of movement. The older participants showed less variability than the young participants (ShAbd: χ2 (1) = 4.42, *p* = 0.036; WrExt: χ2 (1) = 5.62, *p* = 0.018). The effect of target location was found to be significant on shoulder flexion at all the time points assessed (100ms: χ^2^ (2) = 8.08, *p* = 0.018; PKV: χ^2^ (2) = 9.14, *p* = 0.010; PKV50ms: χ^2^ (2) = 7.42, *p* = 0.024; END: χ^2^ (2) = 9.32, *p* = 0.009), shoulder abduction at the time of peak speed and 50 ms after peak speed (PKV: χ^2^ (2) = 9.30, *p* = 0.010; PKV50ms: χ^2^ (2) = 7.39, *p* = 0.025), shoulder external rotation at all the time points except the beginning (PKV: χ^2^ (2) = 13.51, *p* = 0.001; PKV50ms: χ^2^ (2) = 10.34, *p* = 0.006; END: χ^2^ (2) = 7.97, *p* = 0.019), and three DOFs of wrist at the beginning of movement (WrDev: χ^2^ (2) = 12.46, *p* = 0.002; WrExt: χ^2^ (2) = 6.64, *p* = 0.036; WrRot: χ^2^ (2) = 6.13, *p* = 0.047). A significant interaction between age and target location was only found in the variability of shoulder flexion at the beginning of movement (ShFlx: χ^2^ (2) =6.49, *p* = 0.039) and elbow extension at the end of movement (ElFlx: χ^2^ (2) =6.13, *p* = 0.047).

### 3.3. EMG characteristics

Figure 7 illustrates averaged EMG data from seven muscles and tangential speeds of the fingertip for one young and one older subject reaching to three different targets. EMG and fingertip tangential speed signals are time-aligned to movement onset. In both participants, most of the muscles displayed consistent EMG patterns relative to the onset of the movement and some of them are strongly dependent on target position. The activity level in some of the muscles begins to rise before the onset of the movement and there is a substantial amount of activity in the shoulder stabilizer TRP. The BIC EMG exhibits a more complex pattern of activity that started with a large burst, followed by a slight increase to a period of sustained tonic activity.

**Figure 7.**
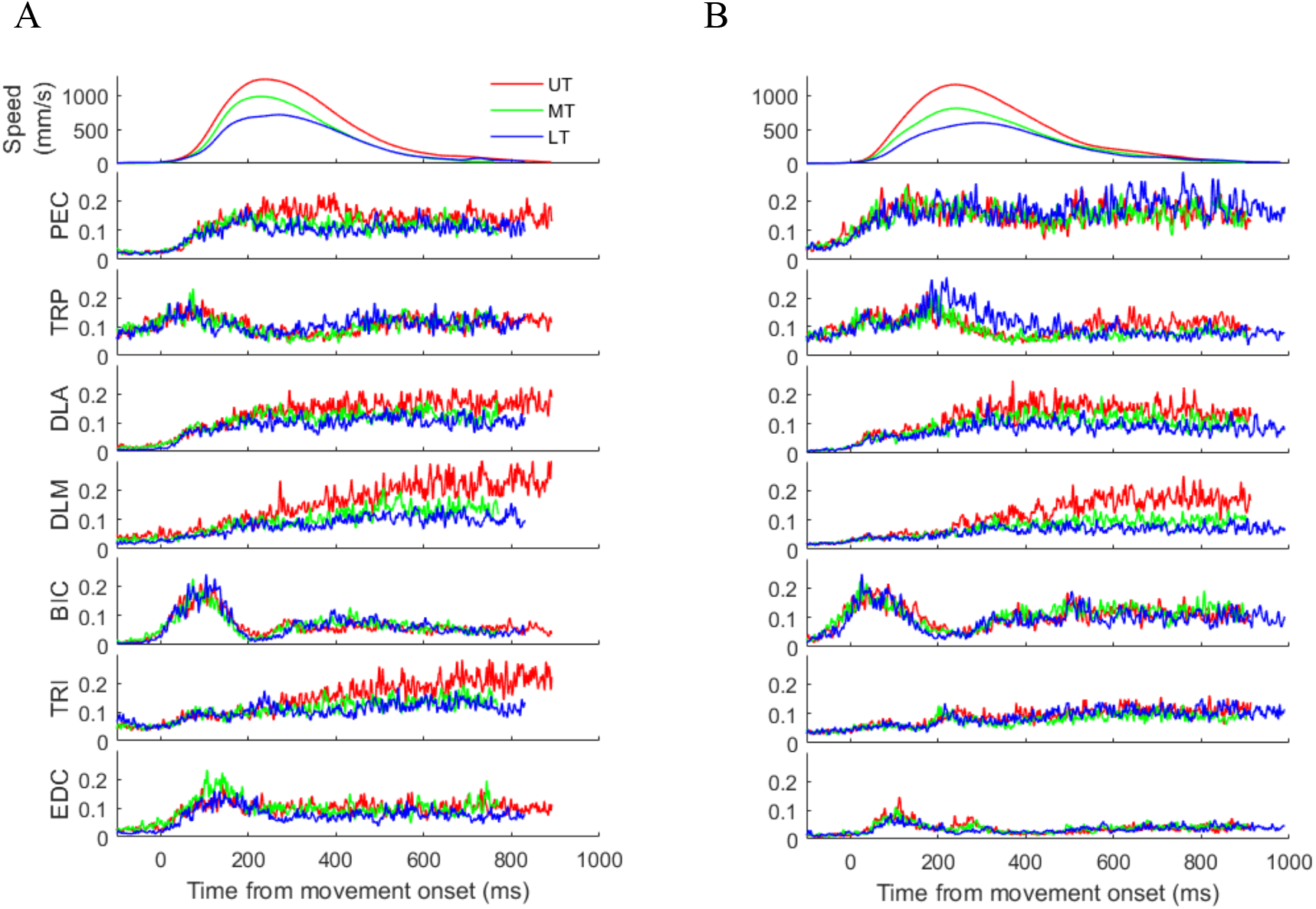
Examples of tangential speed profiles of the fingertip and averaged EMG envelopes from 7 muscles for one young (A) and one older subject (B) reaching to three targets at different heights. EMG signals were normalized with respect to the maximum of the specific muscle over all conditions; muscle abbreviations are defined in the Methods section. Data are aligned to movement onset (Time = 0 ms).

#### EMG onsets

Figure 8 shows mean EMG onsets as a function of target location for all muscles in both young and older groups. The mean onsets of BIC, DLA and TRP (except for LT in older group) were prior to the onset of hand movement. No significant main effect of age group was found in any muscles (all *p* > 0.299). No significant main effect of target height was found in any (all *p* > 0.054) muscles except the TRP (χ^2^ (2) = 9.15, *p* = 0.010); the onset of TRP became progressively later as target height was lowered. No significant interaction between age and target location was found for all muscles, although PEC *(p* = 0.052) and EDC *(p* = 0.056) showed trend.

**Figure 8.**
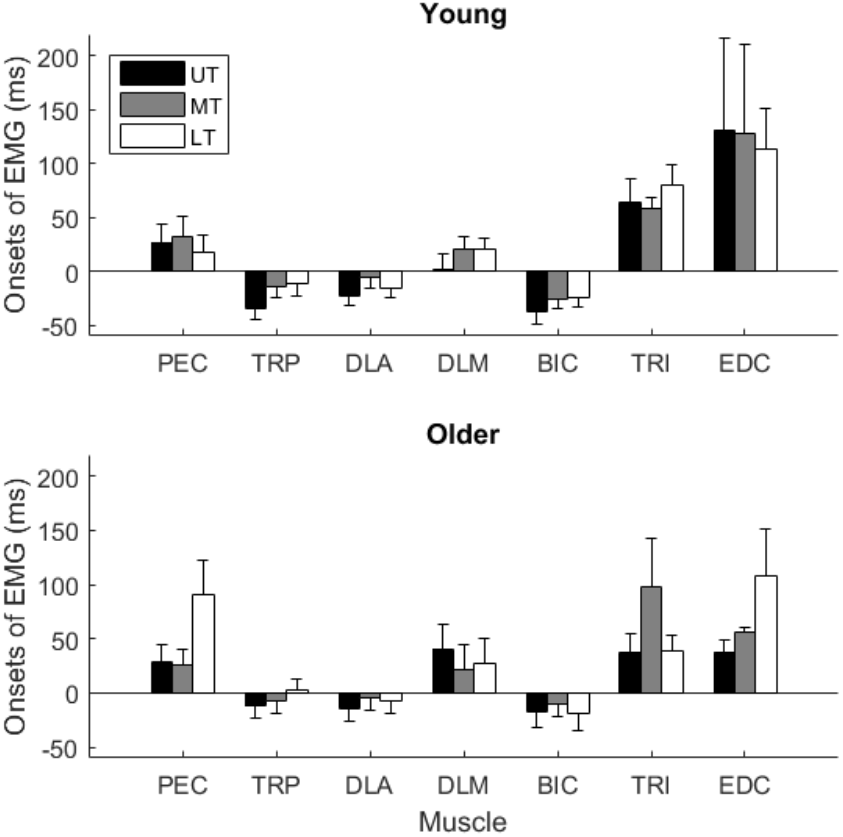
Onsets of EMG plotted as a function of target location for 7 muscles in the young and older subjects (means ± SE). EMG onsets are relative to the onset of hand movement. Negative values indicate onset before the movement onset.

#### EMG peak amplitude

Figure 9 shows mean peak amplitude of normalized EMG as a function of target location for all muscles in both young and older groups. A significant main effect of age was found in muscles TRP (χ^2^ (1) = 4.42, *p* = 0.035), DLA (χ^2^ (1) = 6.01, *p* = 0.014) and BIC (χ^2^ (1) = 5.14, *p* = 0.023). In these three muscles, the older participants had higher peak amplitude than the young participants. Significant main effect of target height was found in muscles DLA (χ^2^ (2) = 29.67, *p* < 0.001), DLM (χ^2^ (2) = 31.70, *p* < 0.001) and TRI (χ^2^ (2) = 29.29, *p* < 0.001). For DLA, pairwise comparisons showed that EMG peak amplitude for three target heights were significantly different from each other. This indicates that the DLA peak amplitude increased with the target height. For both DLM and TRI, pairwise comparisons revealed that EMG peak amplitude to UT were higher than MT (all *p* < 0.001) and LT (all *p* < 0.001), but there was no significant difference between MT and LT (all *p* > 0.307). Neither effect of age (all *p* > 0.317) nor effect of target location (all *p* > 0.395) was found in muscles PEC and EDC. No significant interaction between age and target location was found in any muscles.

**Figure 9.**
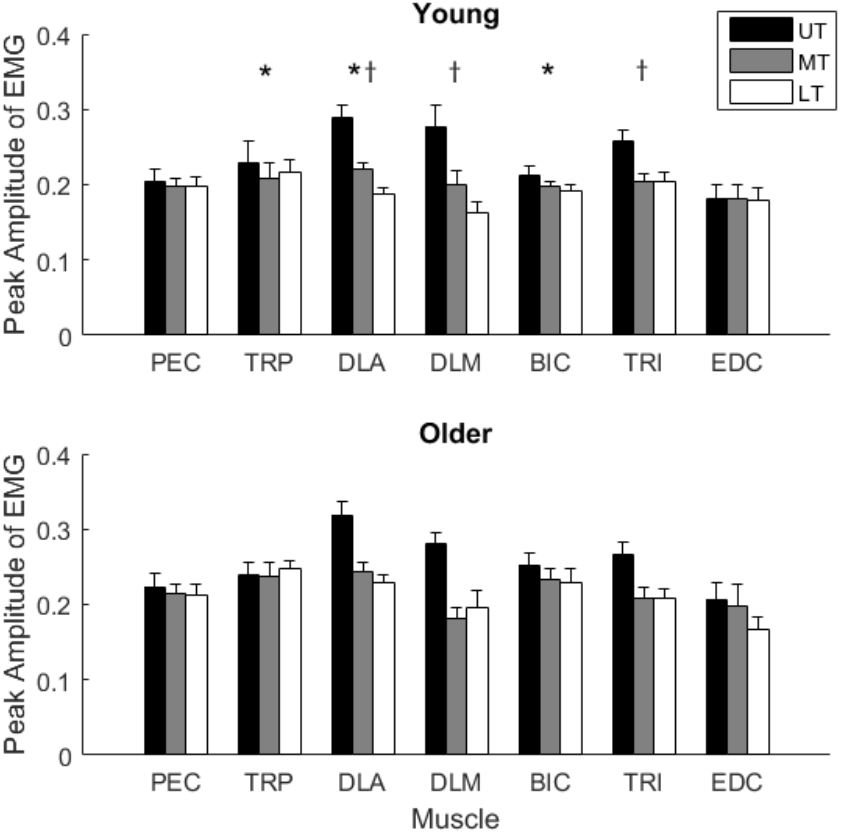
Peak amplitude of normalized EMG plotted as a function of target location for 7 muscles in the young and older subjects (means ± SE). Significant age group differences are indicated by *; target position differences by †.

#### EMG tonic levels at movement end

Figure 10 shows mean tonic activity of normalized EMG as a function of target location for all muscles in both young and older groups. A significant main effect of age was found in muscles BIC (χ^2^ (1) = 9.06, *p* = 0.011) and TRI (χ^2^ (1) = 4.33, *p* = 0.037). In BIC and TRI, the older participants had higher tonic EMG than the young participants. A significant main effect of target height was found in muscles DLA (χ^2^ (2) = 36.25, *p* < 0.001), DLM (χ^2^ (2) = 31.99, *p* < 0.001), TRI (χ^2^ (2) = 30.45, *p* < 0.001) and EDC (χ^2^ (2) = 7.78, *p* = 0.020). For DLA and DLM, pairwise comparisons showed that tonic EMG for three target heights were highly significantly different from each other. This indicates that the tonic activity of DLA and DLM increased with the target height. As the end position is maintained, deltoid activity remains elevated as required to counteract the force of gravity acting on the upper arm. For TRI, pairwise comparisons revealed that tonic EMG to UT were higher than MT *(p* < 0.001) and LT *(p* < 0.001), but there was no significant difference between MT and LT *(p* = 0.388). For EDC, the only difference in tonic EMG was found between UT and LT *(p* = 0.009). Neither effect of age (all *p >* 0.342) nor effect of target location (all *p* > 0.071) was found in muscles PEC and TRP. No significant interaction between age and target location was found in any muscle besides BIC (χ^2^ (2) = 9.06, *p* = 0.011). This suggests that target height had a greater effect on the tonic EMG in biceps muscle in older participants than the young participants.

**Figure 10.**
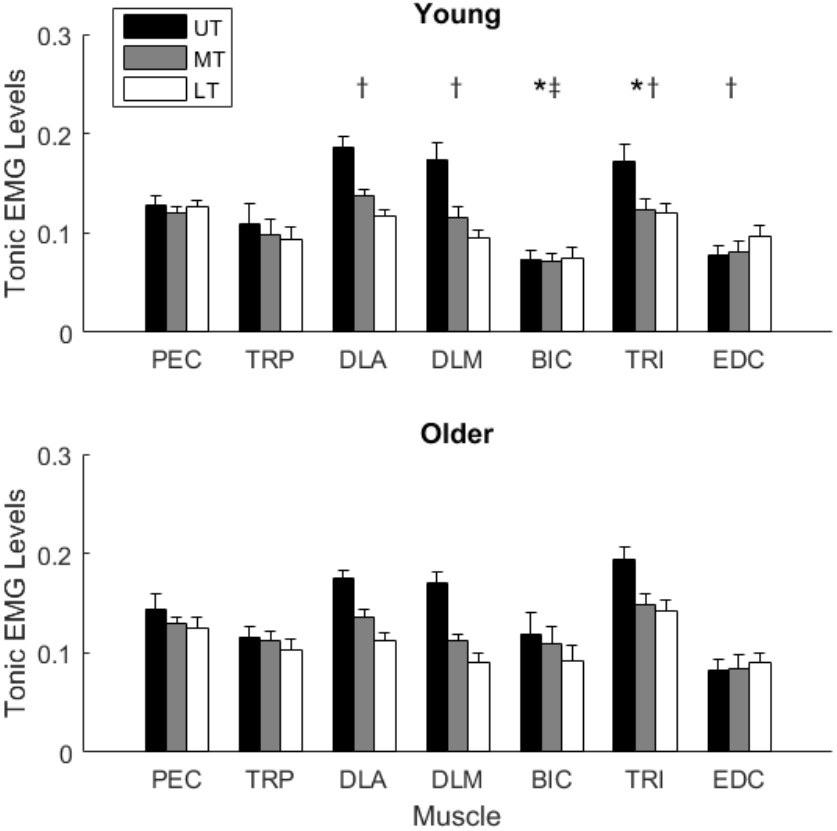
Normalized tonic EMG levels plotted as a function of target location for 7 muscles in the young and older subjects (means ± SE). Significant age group differences are indicated by *; target position differences by †; interaction between age and target position by ‡.

#### Co-contraction of biceps and triceps

Figure 11 shows the mean co-contraction of shoulder-elbow muscles BIC and TRI following reaching movement to the three different targets in young and older groups. There was a significant main effect for age (χ^2^ (1) = 3.89, *p* = 0.048), indicating that the older participants produced a higher level of co-contraction while holding the arm at the final position than the young participants. There was also a significant main effect for target location (χ^2^ (2) = 24.62, *p* < 0.001). Pairwise comparisons revealed that cocontraction following movement to UT were higher than MT and LT (all *p* < 0.001), but no difference between MT and LT *(p* = 0.329).

**Figure 11.**
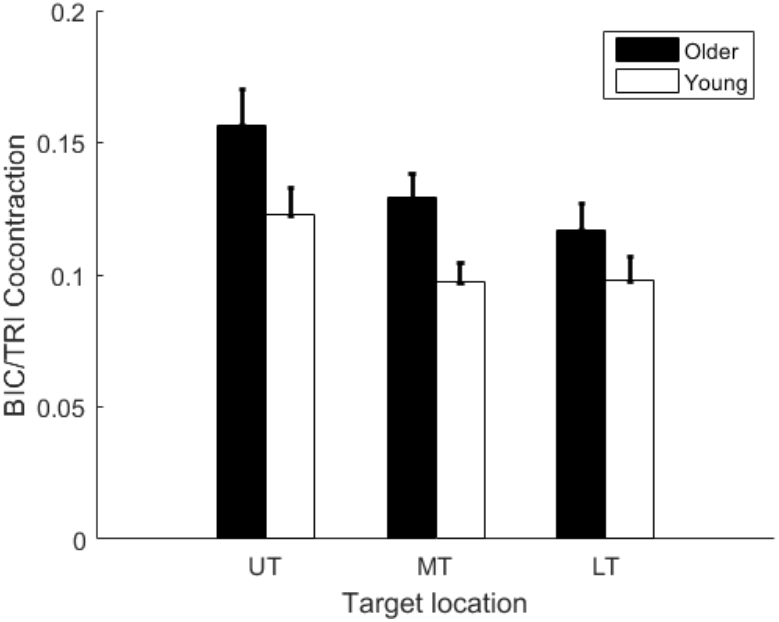
Co-contraction of biceps (BIC) and triceps (TRI) for the three different target locations in the young and older subjects (means ± SE).

To find out whether the co-contraction of BIC and TRI is correlated with the endpoint variability, we plotted the co-contraction after movement end as a function of endpoint variability along the vertical (Z) axis (Figure 12). There was a positive correlation between co-contraction and endpoint variability in the older participants (*r* = 0.47, *p* = 0.019), but not in the young participants (*r* = −0.10, *p* = 0.612). We also tested to see if the co-contraction was correlated with the endpoint variability on the anteroposterior (X) axis. Neither the young (*r* = −0.13, *p* = 0.483) nor the older participants (*r* = 0.28, *p* = 0.185) showed significant correlations.

**Figure 12.**
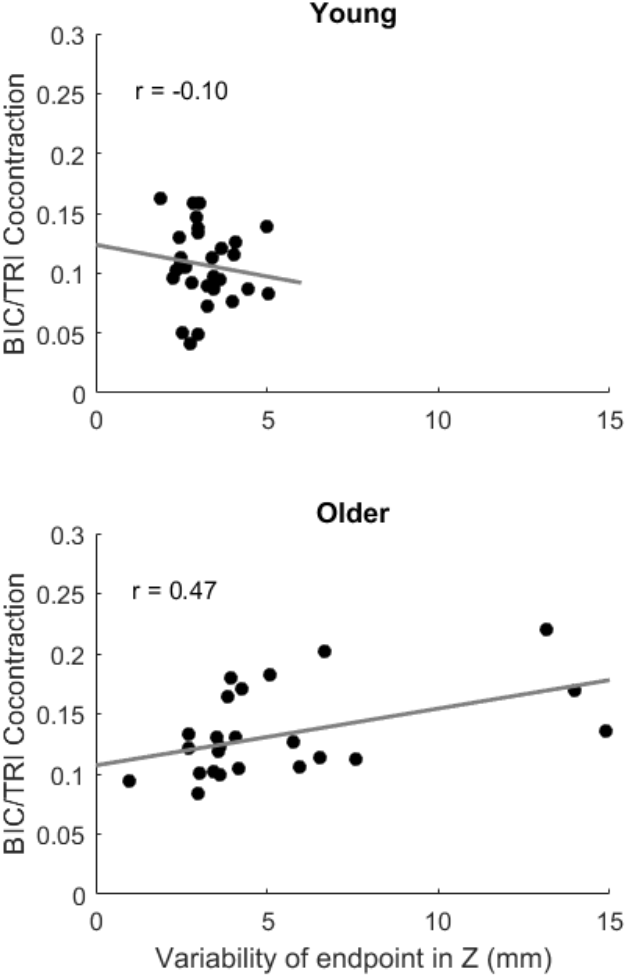
Relationship between co-contraction after movement end and endpoint variability. Mean co-contraction of biceps (BIC) and triceps (TRI) after movement end is plotted as a function of endpoint variability along the vertical (Z) axis. Data for three target locations are pooled for 10 young subjects and 8 older subjects. The gray lines represent the regression lines. The r-value is the correlation coefficient.

## 4. Discussion

In this study, we collected kinematic and electromyographic data to investigate the extent to which aging and vertical target location influence unrestrained reaching movements of the dominant arm against gravity. The emphasis on accuracy rather than speed allowed a closer representation of natural movements made during everyday life. Results showed slower reaction time and greater endpoint variability on the vertical axis in older, as compared to young adults, but reaching performance was similar otherwise. Older participants had more biceps/triceps co-contraction, and co-contraction was related to vertical variability, with more co-contraction correlating with increased variability. They also had higher peak amplitudes in shoulder and elbow muscles.

### Movements against gravity

Most of the variability in endpoint in the older group occurred in the vertical axis, suggesting a poorer prediction or compensation for the effects of gravity. This result extends that of Kimura *et al.* (2015) into a more naturalistic task (i.e., without wrist or finger splints) and demonstrates a deficit in aging with the final corrective movement even with a predicable force such as gravity. Another possibility is a deficit in the previously described elevation/distance *channel* of the sensorimotor transformation (Flanders and Soechting, 1990) as we also found age-related inaccuracy in movement extent. Therefore a single neural control mechanism for both elevation and distance may be impaired with age.

### Hand position and the lower target

The configuration of the target board meant that it was more challenging to reach the lowest target without encountering the horizontal surface with the hand. This may explain the greater variability of the path curvature and endpoint in the x dimension (depth), as the finger/hand orientation could have varied more in order to satisfy this constraint. All participants had less accuracy in movement extent, longer movement times, and more wrist movement for the lower target, among other significant differences for movement to this target as compared to the others. There was no more variability in the vertical dimension, but again, this may reflect the steric constraint of the task.

### Timing

In contrast to the Poston (2009;2013) results, which were with movements performed as fast as possible, there were no differences associated with age in the general structure of reaching movements such as in smoothness or skewness. EMG activation was also similar between the two age groups, except that timing of triceps activation that was affected more by target height in the older group.

### Corrective movements and movement extent

Even with the relatively slow movements we used, there was evidence that movement extent was impaired with age, with errors in the depth dimension, just as was shown in fast movements (Poston et al., 2013). These errors appear to occur later in the movements, as movement extent at the time of peak speed is similar between groups. Gordon (1994) suggested that direction and extent are specified independently in a hand-centered coordinated system. Aging appears to impair the corrective movement extent, even in the context of a non-varying extent requirement. Even though target board was in a fixed position and provided tactile feedback, the older group did not decelerate as accurately as the young group, confirming a deficit in secondary movement planning or execution. And despite this lack of accuracy, there were no differences in movement skewness, suggesting problems with feedback of error. All participants did undershoot the physical target as they braked their movement (Lyons et al., 2006).

Sarlegna (2006) demonstrated longer time to first compensatory movement for a displacement in target, whereas our study involved no target jumps. With the reduced demands of time and compensation, our older group had indistinguishable *temporal* profiles to their movements. This is consistent with the work of (Helsen et al., 2016) that demonstrated good compensation for proprioceptive deficits, when given enough time, and, as in this case, a lack of challenges such as change in target location.

### Kinematic variability

The variability of the reach vector magnitude and variability of the reach vector deviation angle progressed differently during the course of a movement. These results suggest that online control of direction occurs earlier in the movement than the control of movement extent. Our results also indicated that aging impairs the precision of control, around the time of peak speed, of direction but not extent. There were some subtle differences in use of the multiple degrees of freedom of upper extremity joint movement, with less variability in shoulder abduction in the older group and a complex difference in use of shoulder flexion and elbow variability with different targets at different stages of movement. Overall, we were not able to explain differences in endpoint control with differences in individual joint angle variability.

### Muscle activation

The overall pattern of muscle activation was similar between the groups, with some difference in timing and level of activity. The initial burst of biceps activity agrees well with the forces initially required to counteract the force of gravity. The subsequent activity permits controlled, passive extension of the forearm under the force of gravity. The peak levels of activation of key muscles – trapezius, anterior deltoid, and biceps – during each reach (normalized to peak activation across reaches) were higher in the older group. Tonic activation of biceps and triceps was higher in the older group, and target height had more effect on biceps tonic activation in the older group. Our measure of cocontraction of biceps and triceps was predictably higher in the older group. Higher cocontraction was correlated with lower endpoint precision in older adults(Gribble et al., 2003). Taken together, these results suggest that aging affects reaching against gravity by increasing co-contraction as a strategy to compensate for reduced vertical precision. It is also possible that co-contraction is related to decreased strength, as corrective movement speed is decreased. While one would think of co-contraction, with resulting increased impedance, as a strategy to resist unpredictable forces, gravitational forces do change with the changing limb configuration, and prediction of those forces could be less accurate or delayed.

### Limitations and caution in interpretation

#### General issues

As this was a pilot study, there are the usual limitations in interpretation based on numbers of participants. The limitation in numbers also limited the analysis strategy to a between-group comparison, rather than the use of age as a covariate. While all participants were free of any neurological diagnoses, we observed different levels of fitness and ability to follow instructions in the study. These differences were noted but not quantified. We also asked about habitual activities, such as playing musical instruments and other recreational activities that could affect motor function or general fitness. Almost all participants had such activities but this data was not used to subdivide the small sample.

#### Shoulder-Elbow strategy and age

Vandenberghe and colleagues (Vandenberghe et al., 2010) were concerned with relative contributions and timing of shoulder and elbow kinematics in vertical reaching, with a conclusion that shoulder movement led elbow movement, which we also found in examination of joint angles (Fig. 2.) As in that study, we also considered coordination of muscle activity at all times after reach initiation. Our statistical analysis of joint angles was limited to joint excursion and variability. That demonstrated less use of external rotation and more flexion of the shoulder in the older group, as well as less variability of shoulder abduction at the end of the movement. Shoulder flexion variability was affected by age and target location at the beginning of movement and elbow extension at the end of movement, again consistent with the shoulder-elbow strategy. But the EMG analysis showed significant age-dependent differences in proximal *and* distal muscles, and a key role of timing of activation in biceps and triceps, both multijoint (shoulder/elbow) muscles. Normalization of EMG was based on assessment of maximum activity during the reaches, and as with all such studies, normalization could have introduced systematic errors. However, we considered normalization as essential to control for inter-individual differences in body geometry.

Eliminating constraint on the wrist in our study did not have much effect on the qualitative aspects of shoulder and elbow activity during forward reaches, but wrist movement was generally less variable in the older group, suggesting that fewer degrees of freedom were used in control of the arm. (We did not analyze interjoint coordination, particularly because the task timing would not be ideal for that purpose, but that could be done in the future.)

Visual feedback during the reach was reduced due to dim lighting in the room and the lack of illumination of the target LED. However, it was not completely absent, as in some studies. We had previously demonstrated the visual feedback can impair accuracy of arm movements particularly in older people, but in a very different type of task (Boisgontier et al., 2014). Future studies of this sort should use either continuous visual feedback of hand and target or non-visually guided movements to remembered targets.

#### Future directions

We have already acquired data on TMS-induced changes in EMG activity and kinematics for the same participants. The analysis methods and results will allow us to analyze this data and compare the timing and role of frontal lobe areas involved in the control of reaching movements against gravity, as affected by age. (It has been demonstrated that brain activity related to motor performance is more extensive with increasing age (Heuninckx et al., 2005;Heuninckx et al., 2008). This will set the stage for studies of stroke-affected individuals and provide a control for age-related effects, allowing more specific isolation of the effects of focal brain lesions on the cortical motor system. It will also be possible to expand the number of regions tested and type of TMS perturbations in future studies of naturalistic behavior in any study population.

## Conclusions

When unconstrained by speed-accuracy tradeoffs or reduced degrees of freedom of the upper extremity, older adults make kinematically similar reaching movements as younger adults, but with reduced vertical and forward precision and increased co-contraction. Joint co-contraction appears to improve accuracy in the older group, possibly by compensating for unpredictable muscle forces or poor modeling of gravity effects on arm position. There are some quantitative differences in contribution of the shoulder and elbow joints to forward reaching that could be related to changes in muscle mass or joint stiffness.

## Glossary

**Muscle names**
PEC: Pectoralis superior
TRP: Trapezius pars descendens
DLA: Anterior deltoid
DLM: Medial deltoid
DLP: Posterior deltoid
BIC: Biceps brachii
TRI: Triceps long head
FCU: flexor carpi ulnaris
EDC: Extensor digitorum communis
TMS: Transcranial magnetic stimulation
EMG: Electromyelogram
UT: Upper target
MT: Middle target
LT: Lower target

## Author Contributions

Experimental conception and design: GFW, IJ, SPS, OL. Experimental conduct: GFW, NK, LW, FVH, MPB. Data analysis: JT, OL, NK, LW, GFW. Manuscript preparation: GFW, JT. All authors approved the final version of the manuscript.

## References

(1964). Human Experimentation: Code of Ethics of the World Medical Association (Declaration of Helsinki). Can Med Assoc J 91, 619.

Bastiaens, H., Alders, G., Feys, P., Notelaers, S., Coninx, K., Kerkhofs, L., Truyens, V., Geers, R., and Goedhart, A. (2011). Facilitating robot-assisted training in MS patients with arm paresis: a procedure to individually determine gravity compensation. IEEE Int Conf Rehabil Robot 2011, 5975507.

Boisgontier, M.P., and Cheval, B. (2016). The anova to mixed model transition. Neurosci Biobehav Rev 68, 1004–1005.

Boisgontier, M.P., and Nougier, V. (2013). Ageing of internal models: from a continuous to an intermittent proprioceptive control of movement. Age (Dordr) 35, 1339–1355.

Boisgontier, M.P., Van Halewyck, F., Corporaal, S.H., Willacker, L., Van Den Bergh, V., Beets, I.A., Levin, O., and Swinnen, S.P. (2014). Vision of the active limb impairs bimanual motor tracking in young and older adults. Front Aging Neurosci 6, 320.

Bosch, T., De Looze, M.P., Kingma, I., Visser, B., and Van Dieen, J.H. (2009). Electromyographical manifestations of muscle fatigue during different levels of simulated light manual assembly work. J Electromyogr Kinesiol 19, e246–256.

Coats, R.O., Fath, A.J., Astill, S.L., and Wann, J.P. (2016). Eye and hand movement strategies in older adults during a complex reaching task. Exp Brain Res 234, 533–547.

Elliott, D., Hansen, S., Grierson, L.E., Lyons, J., Bennett, S.J., and Hayes, S.J. (2010). Goal-directed aiming: two components but multiple processes. Psychol Bull 136, 1023–1044.

Flanders, M., and Soechting, J.F. (1990). Parcellation of sensorimotor transformations for arm movements. J.Neurosci. 10, 2420–2427.

Goble, D., Coxon, J., Van Impe, A., De Vos, J., Wenderoth, N., and Swinnen, S. (2010). The neural control of bimanual movements in the elderly: Brain regions exhibiting age-related increases in activity, frequency-induced neural modulation, and task-specific compensatory recruitment. Hum Brain Mapp 31, 1281–1295.

Gordon, J., Ghilardi, M.F., Cooper, S.E., and Ghez, C. (1994). Accuracy of planar reaching movements. II. Systematic extent errors resulting from inertial anisotropy. Exp Brain Res 99, 112–130.

Gribble, P.L., Mullin, L.I., Cothros, N., and Mattar, A. (2003). Role of cocontraction in arm movement accuracy. J Neurophysiol 89, 2396–2405.

Grimm, F., Naros, G., and Gharabaghi, A. (2016). Closed-Loop Task Difficulty Adaptation during Virtual Reality Reach-to-Grasp Training Assisted with an Exoskeleton for Stroke Rehabilitation. Front Neurosci 10, 518.

Helsen, W.F., Van Halewyck, F., Levin, O., Boisgontier, M.P., Lavrysen, A., and Elliott, D. (2016). Manual aiming in healthy aging: does proprioceptive acuity make the difference? Age (Dordr) 38, 45.

Heuninckx, S., Wenderoth, N., Debaere, F., Peeters, R., and Swinnen, S.P. (2005). Neural basis of aging: the penetration of cognition into action control. J Neurosci 25, 6787–6796.

Heuninckx, S., Wenderoth, N., and Swinnen, S.P. (2008). Systems neuroplasticity in the aging brain: recruiting additional neural resources for successful motor performance in elderly persons. J Neurosci 28, 91–99.

Kimura, D., Kadota, K., and Kinoshita, H. (2015). The impact of aging on the spatial accuracy of quick corrective arm movements in response to sudden target displacement during reaching. Front Aging Neurosci 7, 182.

Lyons, J., Hansen, S., Hurding, S., and Elliott, D. (2006). Optimizing rapid aiming behaviour: Movement kinematics depend on the cost of corrective modifications. Exp Brain Res 174, 95–100.

Moubarak, S., Pham, M.T., Moreau, R., and Redarce, T. (2010). Gravity compensation of an upper extremity exoskeleton. Conf Proc IEEE Eng Med Biol Soc 2010, 4489–4493.

Moulias, R., Meaume, S., and Raynaud-Simon, A. (1999). Sarcopenia, hypermetabolism, and aging. Z Gerontol Geriatr 32, 425–432.

Oldfield, R.C. (1971). The assessment and analysis of handedness: the Edinburgh inventory. Neuropsychologia 9, 97–113.

Poston, B., Van Gemmert, A.W., Barduson, B., and Stelmach, G.E. (2009). Movement structure in young and elderly adults during goal-directed movements of the left and right arm. Brain Cogn 69, 30–38.

Poston, B., Van Gemmert, A.W., Sharma, S., Chakrabarti, S., Zavaremi, S.H., and Stelmach, G. (2013). Movement trajectory smoothness is not associated with the endpoint accuracy of rapid multi-joint arm movements in young and older adults. Acta Psychol (Amst) 143, 157–167.

Prange, G.B., Jannink, M.J., Stienen, A.H., Van Der Kooij, H., Ijzerman, M.J., and Hermens, H.J. (2009). Influence of gravity compensation on muscle activation patterns during different temporal phases of arm movements of stroke patients. Neurorehabil Neural Repair 23, 478–485.

Prior, S.J., Ryan, A.S., Blumenthal, J.B., Watson, J.M., Katzel, L.I., and Goldberg, A.P. (2016). Sarcopenia Is Associated With Lower Skeletal Muscle Capillarization and Exercise Capacity in Older Adults. J Gerontol A Biol Sci Med Sci 71, 1096–1101.

Przybyla, A., Haaland, K.Y., Bagesteiro, L.B., and Sainburg, R.L. (2011). Motor asymmetry reduction in older adults. Neurosci Lett 489, 99–104.

Sarlegna, F.R. (2006). Impairment of online control of reaching movements with aging: a double-step study. Neurosci Lett 403, 309–314.

Vandenberghe, A., Levin, O., De Schutter, J., Swinnen, S., and Jonkers, I. (2010). Threedimensional reaching tasks: effect of reaching height and width on upper limb kinematics and muscle activity. Gait Posture 32, 500–507.

Vandervoort, A.A. (2002). Aging of the human neuromuscular system. Muscle Nerve 25, 17–25.

Welsh, T.N., Higgins, L., and Elliott, D. (2007). Are there age-related differences in learning to optimize speed, accuracy, and energy expenditure? Hum Mov Sci 26, 892–912.

